# A novel method (RIM-Deep) for enhancing imaging depth and resolution stability of deep cleared tissue in inverted confocal microscopy

**DOI:** 10.1101/2024.07.19.604108

**Authors:** Yisi Liu, Pu Wang, Junjie Zou, Hongwei Zhou

## Abstract

The increasing use of tissue clearing techniques underscores the urgent need for cost-effective and simplified deep imaging methods. While traditional inverted confocal microscopes excel in high-resolution imaging of tissue sections and cultured cells, they face limitations in deep imaging of cleared tissues due to refractive index mismatches between the immersion media of objectives and sample container. To overcome these challenges, the RIM-Deep was developed to significantly improve deep imaging capabilities without compromising the normal function of the confocal microscope. This system facilitates deep immunofluorescence imaging of the prefrontal cortex in cleared macaque tissue, extending imaging depth from 2 mm to 5 mm. Applied to an intact and cleared Thy1-EGFP mouse brain, the system allowed for clear axonal visualization at high imaging depth. Moreover, this advancement enables large-scale, deep 3D imaging of intact tissues. In principle, this concept can be extended to any imaging modality, including existing inverted wide-field, confocal, and two-photon microscopy. This would significantly upgrade traditional laboratory configurations and facilitate the study of connectomics in the brain and other tissues.

## Introduction

Tissue clearing techniques enable 3D imaging of whole tissues and organs[1]. Unlike traditional histology, these techniques reduce light scattering and absorption, enhancing depth imaging capabilities and achieving single-cell resolution in deep samples [2]. These approaches is now widely used in neurobiology, developmental biology, immunology, and oncology. To achieve deep imaging of fully cleared tissues, various optical sectioning techniques for three-dimensional imaging have rapidly developed, including light sheet microscopy, two-photon microscopy, and confocal microscopy[1].

For effective 3D imaging of cleared tissues, it is crucial to utilize microscopes equipped with objectives that offer both low to medium magnification and long working distances [3]. Objectives with low numerical aperture (NA) are often used to achieve these long working distances, but they face significant challenges in maintaining high axial resolution for deep tissue imaging [4]. The main issue arises from light scattering and reflection at the interfaces within the sample and between the sample and the lens, leading to image distortion and blurring. This phenomenon, known as spherical aberration, occurs when light rays entering the periphery of the lens focus at a different point on the optical axis compared to those entering near the center, leading to a significant degradation in fluorescence signal intensity and image resolution [5]. Using dry objectives with high-refractive-index mounting media or cleared tissues can exacerbate this issue, as the sample itself contributes to the aberration [6]. Therefore, matching the refractive index (RI) of the objective’s immersion medium to that of the sample’s immersion medium is essential to mitigate axial distortion caused by spherical aberration.

Many immersion objectives feature correction collars that can be adjusted to compensate for RI mismatches. Achieving an exact match between the refractive indices of the immersion media and the sample medium is often challenging. Although dipping caps with glass windows can transform air objectives into immersion lenses by enabling the front lens to be directly inserted into the sample media, this approach helps to avoid axial deviation during imaging. However, immersion objectives on inverted microscopes encounter limitations [7]. These objectives cannot be inserted into sample chambers and must move up and down outside the chamber, which restricts the imaging depth.

To overcome these limitations, several alternative methods have been developed. One approach involves moving the sample within a chamber filled with the mounting media while keeping the imaging objective fixed outside the chamber[8]. Another technique utilizes adaptive optics, such as deformable mirrors or spatial light modulators, to correct aberrations across the field of view [9]. Additionally, radially symmetric phase masks are used to balance spherical aberration at various depths within the sample, thereby simplifying the image restoration process through post-capture deconvolution [10].

However, inverted confocal microscopes encounter issues due to the gravitational dispersion of the RI buffer between the objective and the confocal dish, resulting in RI mismatches with air and complicating high-resolution imaging. Additionally, the use of long-working-distance air objectives causes severe spherical aberrations due to RI mismatches, further limiting imaging depth and resolution.

Here, we developed the Refractive Index Matching-Deep (RIM-Deep) system to address these challenges, which includes an immersion chamber designed around the optical components of the objective lens and a specimen holder integrated with a motorized x–y–z stage. This design effectively stabilizes the refractive index between the objective and the sample media. When paired with an immersion objective, the RIM-Deep system enables high-resolution, deep imaging, significantly enhancing the capabilities of inverted confocal microscopy. This innovation provides a practical solution for achieving high-quality 3D imaging of cleared tissues, paving the way for more detailed and accurate biological studies.

## Materials and Methods

### Animals

We used 6-year-old male Macaca fascicularis brain and male Thy1-EGFP mice brain gifted from Prof. Xiong Cao (South Medical University). All animal procedures were approved by the Southern Medical University Animal Ethics Committee and conducted in compliance with the guidelines of the Chinese Council on Animal Care to minimize animal suffering and reduce the number of animals used.

### Thy1-EGFP mouse brain preparation for clearing

We employed the CUBIC method for clearing Thy1-EGFP mouse fixed brain tissue [11], with the process carried out at the Shared Instrument Platform of the School of Life Sciences at Tsinghua University.

### Vascular labeling and tissue clearing

Adult male C57BL/6J mice (8 weeks) were used for the creation of the photothrombotic stroke model. Mice were anesthetized with sodium pentobarbital (50 mg/kg, IP). The head was secured in a stereotaxic device, and the skull was exposed. A photoactive dye, Rose Bengal was injected into the tail vein. Focused 561 nm laser irradiation was then applied to the cortical area to activate the dye, causing thrombosis and inducing ischemia by obstructing blood vessels. After surgery, mice were placed in warm cages for recovery and subsequently returned to their cages. Three days later, mice underwent deep sedation via intraperitoneal administration of sodium pentobarbital at a dosage of 50 mg/kg. Following this, vascular labeling was executed utilizing the VALID protocol [12]. Initially, mice were transcardially infused with 0.01 M phosphate-buffered saline (PBS) for the removal of blood from the vascular system. Subsequently, a volume of 10–15 mL of VALID working solution, warmed to maintain its fluid state, was perfused through the circulatory system. Post perfusion, the mice were refrigerated at 4°C overnight to solidify the gel within the vasculature. The next phase involved organ extraction and further post-fixation in 4% paraformaldehyde (PFA) for an extended period. Extreme care was taken to detach the skull from the perfused mouse body to avoid distortion of the sample. During this process, the dura mater was carefully excised. Finally, the removed skull was post-fixed in 4% PFA for an overnight duration.

MACS was executed for demonstrated previously[13]. Fixed samples were serially incubated in MACS-R0, MACS-R1, and MACS-R2 solutions, with gentle shaking at room temperature. MACS-R0 was prepared by mixing 20% (vol/vol) MACS with 15% (wt/vol) sorbitol (Sigma, 85529) in dH2O; MACS-R1 was prepared by mixing 40% (vol/vol) MXDA with 30% (wt/vol) sorbitol dissolved in 1× PBS; MACS-R2 was prepared by mixing 40% (vol/vol) MXDA with 50% (wt/vol) sorbitol in dH2O.

### Sample pretreatment and immunolabeling

As previously described [14], The Macaca fascicularis fixed brain underwent a series of washes and treatments as follows: Initially, they were washed twice for one hour each in PBS, followed by sequential hour-long immersions in 50%, 80%, and 100% methanol solutions, with the latter repeated. The samples were then bleached overnight at 4°C using a 5% hydrogen peroxide solution in a 20% DMSO/methanol mix (prepared with 1 volume of 30% H_2_O_2_, 1 volume of DMSO, and 4 volumes of methanol, all kept ice cold). Post-bleaching, the samples were washed twice in methanol for an hour each, followed by two one-hour washes in a 20% DMSO/methanol solution, then in 80% and 50% methanol solutions for an hour each, and finally, twice in PBS for an hour. The last step involved two one-hour washes in a PBS solution containing 0.2% Triton X-100, preparing the samples for subsequent staining procedures.

The samples underwent a series of incubation and washing processes as described: Initially, they were incubated overnight at 37°C in a solution of PBS with 0.2% Triton X-100, 20% DMSO, and 0.3 M glycine. Following this, they were blocked using a PBS solution containing 0.2% Triton X-100, 10% DMSO, and 6% Donkey Serum at 37°C for 3 days. Subsequently, the samples were washed twice for an hour each in a PBS solution with 0.2% Tween-20 and 10 μg/ml heparin (PTwH), then incubated in primary antibody (Anti-CD31, ab281583) dilutions in a PTwH solution with 5% DMSO and 3% Donkey Serum at 37°C for 4 days. After this incubation, the samples were washed in PTwH for one day, followed by an incubation in secondary antibody (Alexa Fluor® 594, ab150080)) dilutions in a PTwH solution with 3% Donkey Serum at 37°C for 4 days. The final step involved washing the samples in PTwH for two days before proceeding with clearing and imaging.

The immunolabeled tissues underwent a clearing process using the iDISCO technique [15], CUBIC or MACS. This method and the solutions employed were adapted from protocols detailed in a previously published study. The entire procedure was conducted at the Shared Instrument Platform located in the School of Life Sciences at Tsinghua University.

### Microscope and objectives

For image acquisition, we used the Nikon AXR inverted laser scanning confocal microscope and Leica STELLARIS 5 inverted laser scanning confocal microscope, paired with a Nikon 10x/0.5 NA immersion objective.

### Design principle of the RIM-Deep

The RI adapter consists of a solution reservoir (Supplementary figure 1A), a specimen holder and support bracket (Supplementary figure 1B). The solution reservoir and specimen holder are made of glass. The nested arrangement of the solution and specimen holder forms a semi-closed space at the outer reservoir, which is filled with imaging buffer (Supplementary figure 1C-F). The specimen holder is centrally positioned on the top surface of the support bracket, ensuring that its base is parallel to the base of the support bracket. The bottom of the specimen holder is 0.17 mm thick cover glasses. The solution reservoir is nested in the cap of the objective.

### Image acquisition

Cleared tissue was placed in a 35 mm confocal dish or the dual-reservoir nested adapter. To ensure that the sample does not slide during X-Y scanning, the sample was fixed in place with transparent glass glue before imaging. Unless otherwise specified, the Z intensity correction module was used to offset the fluorescence signal degradation with depth. The NIS-Elements Advance research software was used for post-processing of images. The processed data were then imported into Imaris (Version 9.0.1, Bitplane AG) for the next step.

## Result

Official adapters for 10X immersion objectives in inverted confocal microscopy significantly enhance cleared tissue imaging, but RI buffer leakage still occurs, causing RI mismatch and limiting imaging depth (Supplementary Figure 2A). In response to this challenge, the RIM-Deep was designed based on a light sheet microscopy imaging chamber. It features a media reservoir affixed to the objective and filled with RI buffer, enabling 3D imaging of the sample within an imaging buffer. To further optimize imaging performance, a support bracket integrated into the motorized stages elevates the imaging platform and centrally positions a specimen holder (Supplementary Figure 1 and 2B).

To demonstrate the effectiveness of the RIM-Deep, imaging resolution was characterized by comparing 3-μm diameter fluorescence beads using standard adapter versus the RIM-Deep in 10X immersion objective (Figure 1A and B). Within the 2mm imaging limit, 10X immersion objectives with the standard adapter maintain both optical axial and lateral resolution. As expected, the RIM-Deep also maintained optimal axial resolution across 5mm. (Figure 1C-E). However, there was no significant difference in lateral (XY) resolution between the two types of chamber (Figure 1C-E).

**Figure 1.**
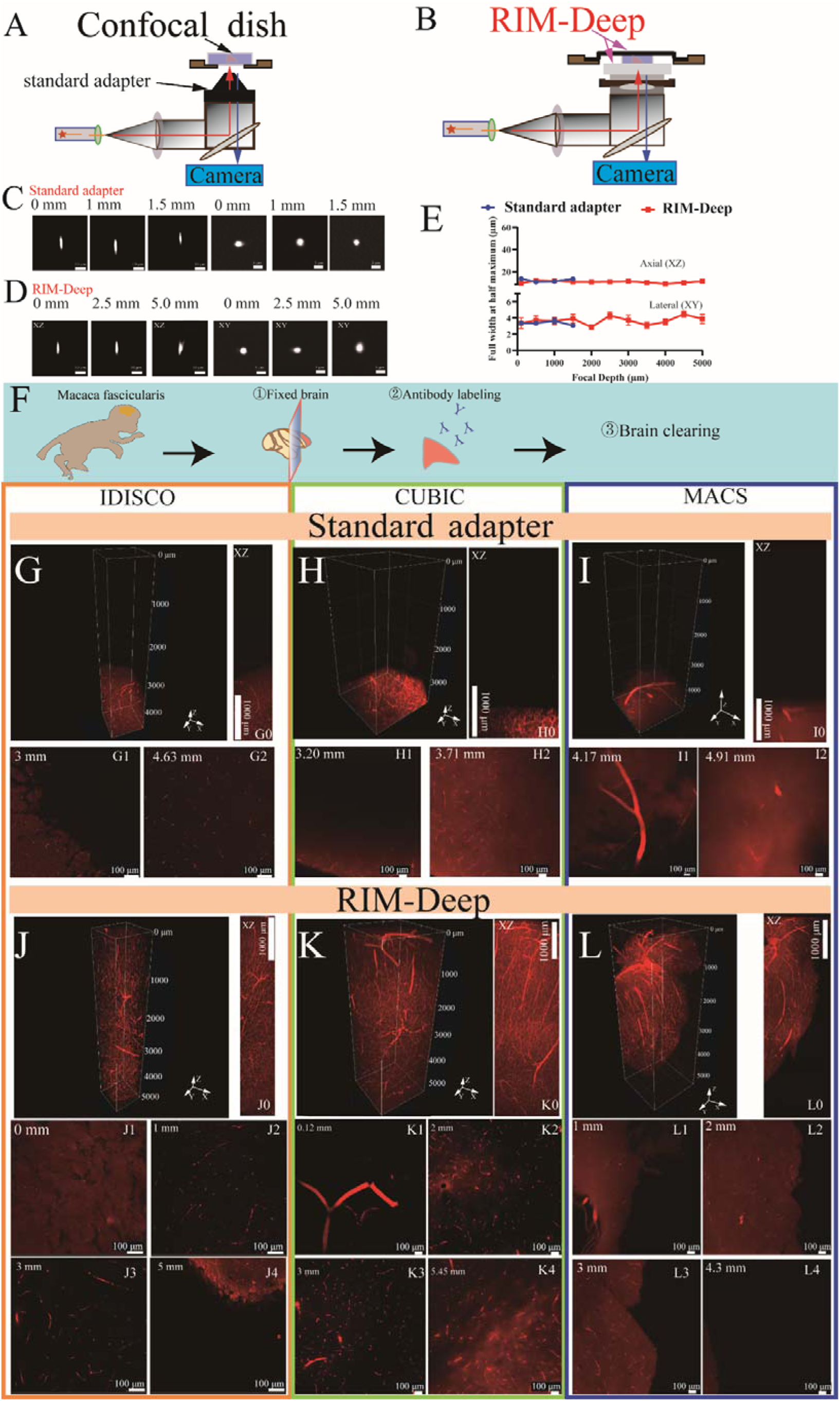
Resolution characterization and image depth of standard adapter and RIM-Deep. (A and B) Schematic diagram of a 10X immersion objective with a standard adapter (A) or RIM-Deep (B). (C-D) MIP of 3-μm-diameter beads imaged in the *xy* and *yz* planes using the standard adapter (C) or RIM-Deep (D) at different axial positions. (E) Axial resolutions for a 10X immersion objective paired with standard adapter or RIM-Deep at different axial positions. The resolution is estimated by FWHMs of intensity profiles with a Gaussian fit for 3-μm-diameter beads embedded in 1% agarose dissolved in CUBIC mounting solution. Data are presented as mean ± s.e.m. (F) The experimental scheme for the brain clearing process in *Macaca fascicularis*. (G, H, I) Three-dimensional reconstruction of the *Macaca fascicularis* brain vasculature using three different tissue clearing methods (iDISCO, CUBIC or MACS) with a standard adapter. (G0, H0, I0) MIP of G, H or I in *xz* plane. (G1-G2, H1-H2, I1-I2) Optical section of (G, H, I) at varying depths. (J, K, L) Three-dimensional reconstruction of the *Macaca fascicularis* brain vasculature using (iDISCO, CUBIC or MACS) with a RIM-Deep. (J0, K0, L0) MIP of J, K or L in *xz* plane. (J1-J4, K1-K4, L1-L4) Optical section of (J, K, L) at varying depths.

Subsequently, the high-depth imaging performance of the RIM-Deep was evaluated and compared to the standard adapter. Imaging was performed on a *Macaca fascicularis* brain section cleared using the iDISCO, CUBIC, and MACS methods (Figure 1F). Gravity affected the refractive index matching fluid at the tip of the objective, limiting the imaging depth to less than 2 mm (Figure 1 G-I2, Rich Media - Video 1 - 3). As observed, with the RIM-Deep, the specimen holder was fully immersed in the RI liquid, the depth of images is obviously enhanced to about 5 mm, and the vessel structures could thereby be observed clearly (Figure 1 J-L4, Rich Media - Video 4 - 6).

To further demonstrate the resolution and depth robustness of the RIM-Deep, experiments were conducted on Thy1-EGFP mouse brains cleared with the CUBIC method (Figure 2A). The use of the RIM-Deep also significantly extended the imaging depth to approximately 5 mm (Figure 2B, Rich Media - Video 7). This extension enabled the clear visualization and reconstruction of neuronal soma and axon structures in the hippocampus and thalamus (Figure 2C, E, F, G, I), maintaining resolution at the micron level (Figure 2D and H).

**Figure 2.**
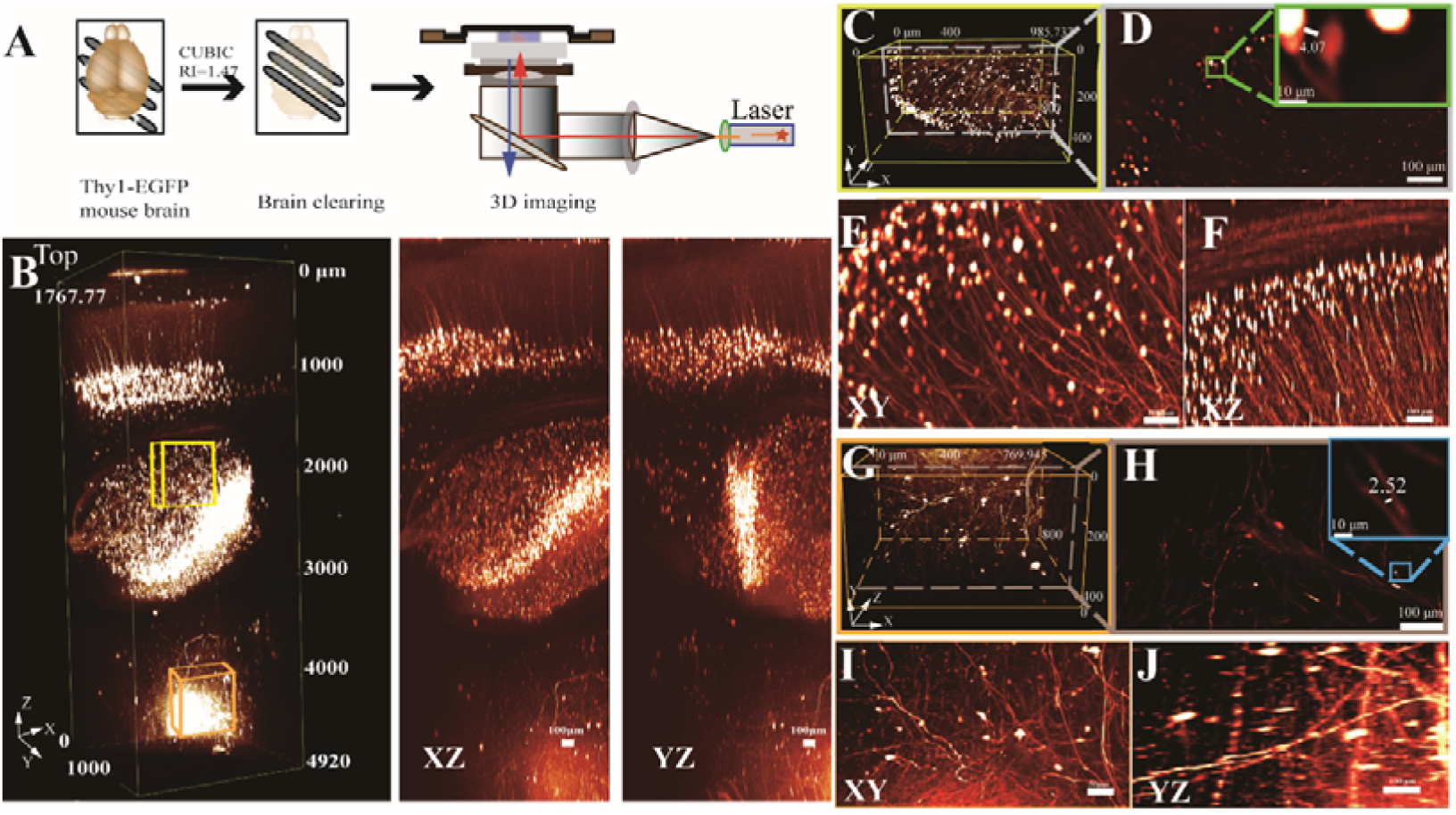
Large-depth imaging of Thy 1-EGFP mouse brains using RIM-Deep. (A) Experimental scheme. (B) 3D reconstruction of the ∼5 mm deep in the mouse brain (left), MIP views in XZ (middle) and YZ (right). (C and G) 3D reconstructions of neuronal structures within the hippocampus (C, yellow box) and thalamus (G, orange box), respectively, as indicated in B. (D and H) Lateral slices through the indicated lateral planes in (C and G). Zoom-in views of the selected areas in top right. (E-F) MIP view of hippocampus(C) in *xy* and *xz*. (I-J) MIP view of thalamus (G) in *xy* and *yz*.

A 3×3 stitching mode with multi-layer Z-axis scanning was used to examine a cleared brain segment (Figure 3A-B, Rich Media - Video 8). This method enabled visualization of neuronal cell bodies and axons up to nearly 5 mm in X-Y optical slices (Figure 3C). For large tissue imaging, the RIM-Deep was used on a cleared half-brain, achieving comprehensive imaging (Figure 3D). Its stable and precise positioning ensured accurate alignment across all optical slices, facilitating detailed and thorough imaging (Figure 3E-I).

**Figure 3.**
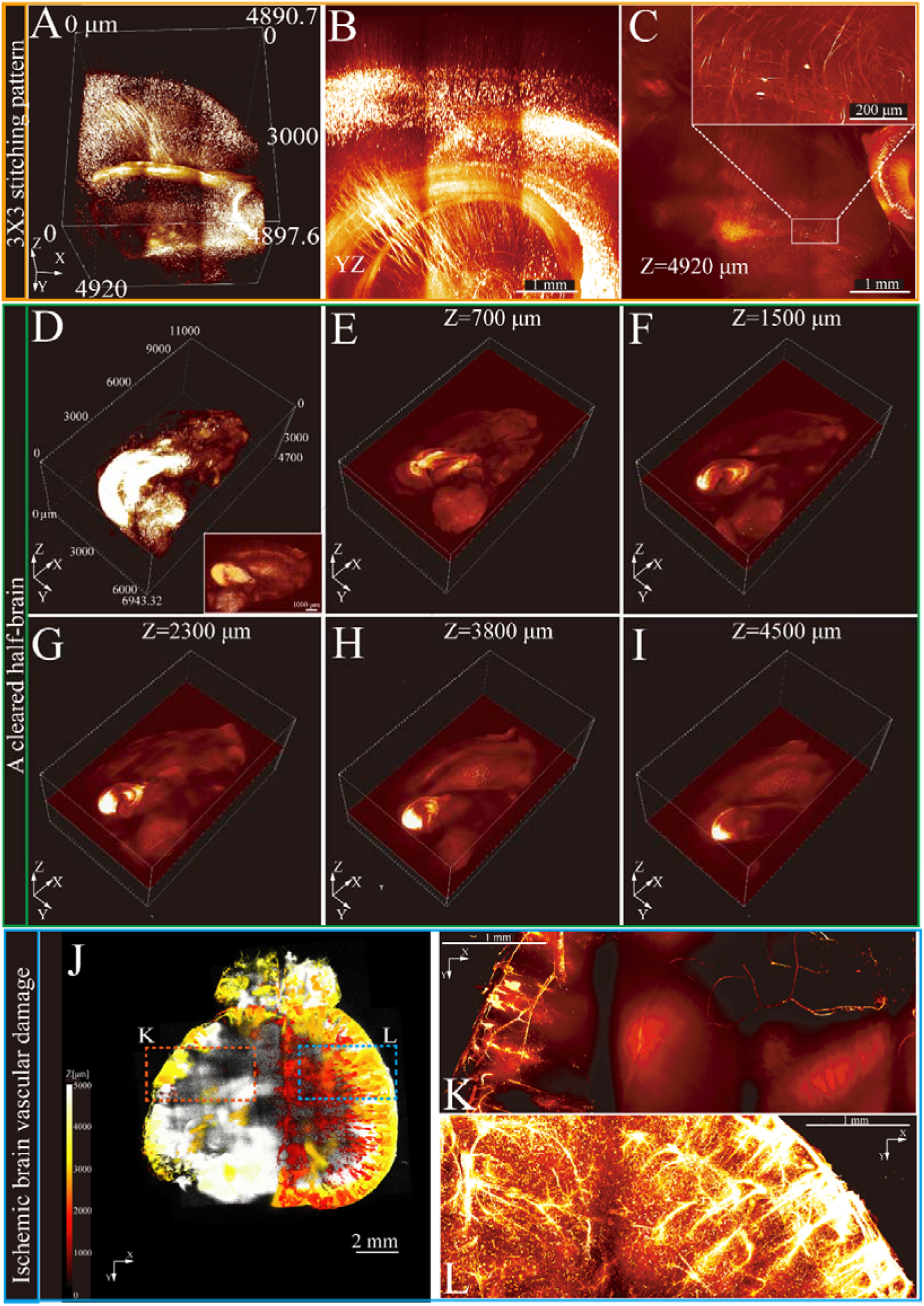
3D imaging and reconstruction of neural and vascular structures in intact brain tissues using RIM-Deep. (A) 3×3 stitching pattern of deep imaging of a cleared brain. (B) MIP of *yz* side view of A. (C) Optical section of top layer in (B). Zoom-in views of the selected areas in top right. (D) 3D reconstruction of a half of cleared brain in a Thy1-eGFP mouse brain. The white box represents MIP. (E-I) Stitched single layer images in the Z-direction. (J) 3D imaging of the entire brain vasculature in ischemic stroke mice. The images along the z stack are colored by spectrum. (K) MIP of vascular imaging in the ischemic region (red box in J). (L) MIP of vascular imaging in the contralateral region (blue box in J).

Stroke, caused by disrupted brain blood supply, induces cell death and significant vascular changes [16]. Generally, confocal microscopy is limited to thin sections and cannot fully study vascular networks. However, with the assistance of the RIM-Deep, complete observation of vascular damage is possible. Therefore, mouse brains with induced cortical ischemia were examined to thoroughly investigate this process. Using VALID labeling and MACS tissue clearing techniques, the cerebral vasculature was reconstructed, revealing a significantly reduced vessel density in the ischemic cortex (Figure 3J-L, Rich Media - Video 9 - 11). These findings demonstrate the effectiveness of the RIM-Deep in visualizing disease-related changes in vascular structures, making it a valuable tool in neuroscience research.

The performance of the RIM-Deep method was evaluated using the Leica inverted confocal microscope. Undoubtedly, high-resolution imaging of multiple cross-sections was achieved for a monkey brain vasculature with a depth approaching 5 mm, demonstrating consistent results across different imaging platforms (Supplementary Figure 3, Rich Media - Video 12).

## Discussion

Inverted confocal microscopy often faces challenges such as limited axial resolution and spherical aberration when imaging cleared tissues. The RIM-Deep, featuring a media reservoir filled with RI buffer, was designed to address these issues. Experimental validations showed that the RIM deep maintained optimal axial resolution when imaging fluorescent beads. Using the 3 different tissue cleaning method on Macaca fascicularis brains, imaging depths reached up to 5 mm with clear visualization of vascular structures. Similarly, in Thy1-eGFP mouse brains, the RIM deep extended imaging depth to approximately 5 mm, allowing clear visualization of neuronal and axonal structures. The RIM deep also effectively visualized vascular damage in mouse brains with cortical ischemia, proving valuable in neuroscience research. The RIM-Deep method also demonstrates versatility across different imaging platforms, ensuring consistent performance in various experimental setups.

The development of tissue clearing has revolutionized neuroscientific research. However, two-photon and light sheet microscopes are prohibitively expensive and their operation and maintenance costs are high, limiting their application scenarios. Many hospitals and laboratories, although capable of performing tissue clearing independently, are unable to conduct imaging. The inverted point-scanning confocal microscope, preferred for live cell and tissue slice imaging, offers advantages in deep imaging of cleared tissues over upright microscopes, including reduced sample contact and buffer use, lowering the risk of damage and cost. However, the use of long-working-distance air objectives introduces severe spherical aberrations due to RI mismatch, limiting imaging depth and resolution. Our RIM-Deep addresses these issues, extending imaging depth without compromising the microscope’s normal functionality.

Wide-field microscopes, although efficient for rapid imaging, often face limitations in depth and resolution when applied to cleared tissues[17]. The RIM-Deep can also be employed in these cases to enhance imaging performance. When using wide-field microscopy with the RIM-Deep, it is necessary to apply deconvolution algorithms or other techniques to eliminate out-of-focus signals[17]. However, it is important to note that this system may reduce fluorescence intensity in deeper regions of the sample.

### Limitations of the study and prospect

Despite the advantages offered by the RIM-Deep, several limitations remain. The imaging speed of inverted point-scanning confocal microscopy is slower compared to light sheet microscopy, necessitating advanced post-processing algorithms for high-speed 3D imaging. To address this, resonance scanning methods combined with artificial intelligence-based image analysis can significantly improve the efficiency and accuracy of imaging, providing a more robust solution for detailed analysis [18]. Iterative reconstruction algorithms can enhance resolution and mitigate imaging artifacts. However, achieving uniform resolution across different depths remains challenging, particularly for very dense samples.

Furthermore, our research primarily focuses on neural tissues, and its effectiveness in imaging other tissue types remains to be explored. What’s more, the objective lens used is limited to a correction collar for refractive indices between 1.33 and 1.51. Techniques like iDISCO and aqueous CUBIC/MACS, with refractive indices of 1.56 and 1.51/1.52 respectively, facilitate depth imaging to nearly 5 mm. However, if the media’s refractive index exceeds the lens’s correction range, it could compromise resolution and image quality. Recently, laboratories have developed objectives with long working distances, high numerical apertures, and broad refractive index correction capabilities [19]. The Schmidt immersion objective, composed of spherical mirrors and aspheric correction plates, achieves an NA of 1.08 at a refractive index of 1.56, FOV of 1.1 mm, and WD of 11 mm [19].

Future work should explore machine learning-based techniques to further optimize image quality and resolution [20]. Moreover, as imaging depth increases, maintaining a high signal-to-noise ratio (SNR) becomes more challenging. Advanced denoising algorithms will be critical to address these issues, ensuring high-quality imaging of thick tissue samples [21, 22].

In summary, the RIM-Deep represents a significant advancement in deep tissue imaging for inverted confocal microscopy, addressing key challenges related to axial resolution and spherical aberration. However, further refinements and integration with advanced computational techniques are necessary to optimize its performance and expand its applicability in various research fields.

## Supporting information

Rich Media - "Figure 1G- Video 1 - Three-dimensional reconstruction of the Macaca fascicularis brain vasculature iDISCO with the standard adapter"

Rich Media - "Figure 1H - Video 2 - Three-dimensional reconstruction of the Macaca fascicularis brain vasculature using CUBIC with the standard adapte

Rich Media - "Figure 1I - Video 3 - Three-dimensional reconstruction of the Macaca fascicularis brain vasculature using MACS with the standard adapter

Rich Media - "Figure 1J- Video 4 - Three-dimensional reconstruction of the Macaca fascicularis brain vasculature iDISCO with RIM-Deep"

Rich Media - "Figure 1K - Video 5 - Three-dimensional reconstruction of the Macaca fascicularis brain vasculature using CUBIC with RIM-Deep."

Rich Media - "Figure 1L - Video 6 - Three-dimensional reconstruction of the Macaca fascicularis brain vasculature using MACS with the RIM-Deep."

Rich Media - "Figure 2 - Video 7 - Deep imaging of a single field in cleared Thy1-EGFP mouse brain tissue"

Rich Media - "Figure 3A-C - Video 8 - Deep imaging in 3×3 tiling mode of cleared Thy1-EGFP mouse brain tissue"

Rich Media - "Figure 3J - Video 9 - Vascular network of the whole brain in MCAO mouse"

Rich Media - "Figure 3K- Video 10 - Vascular network of the ischemic side in MCAO mouse brain"

Rich Media - "Figure 3L - Video 11- Vascular network of the control side in MCAO mouse brain"

Rich Media - "Supplementary figure 3 - Video 12 - imaging of Macaca fascicularis brain vasculature using Leica STELLARIS 5 with RIM-Deep."

## Acknowledgments

This work was supported by grants from the National Key R&D Programme of China (2022YFA0806400), Guangzhou Key Research Programme on Brain Science (202206060001) and the National Natural Science Foundation of China (82130068) to H- WZ.

## Author contributions

Yisi Liu and Pu Wang designed and supervised the study. Yisi Liu collected and analysed the data. Yisi Liu and Junjie Zou collected samples. Yisi Liu wrote the manuscript. Hongwei Zhou contributed to text revision and discussion. All authors discussed the results and approved the manuscript.

---

**Supplementary figure 1.**
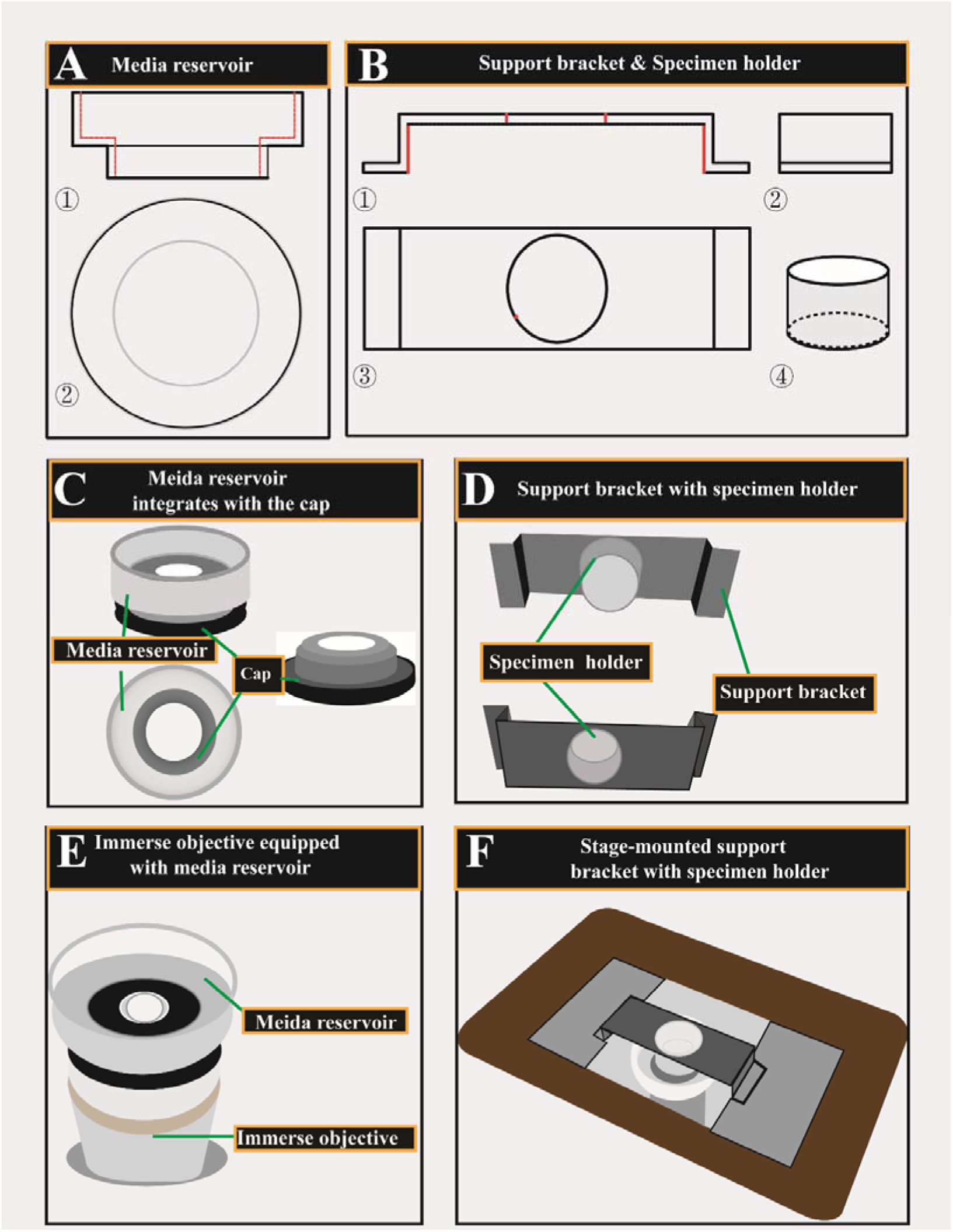
Design and configuration of RIM-Deep for inverted confocal microscope. (A) A Three-view diagram of a media reservoir. (B) ①-③, Three-view diagram of the support bracket. ④, Three-dimensional diagram of the specimen holder. (C-F) RIM-Deep assembly procedure.

**Supplementary Figure 2.**
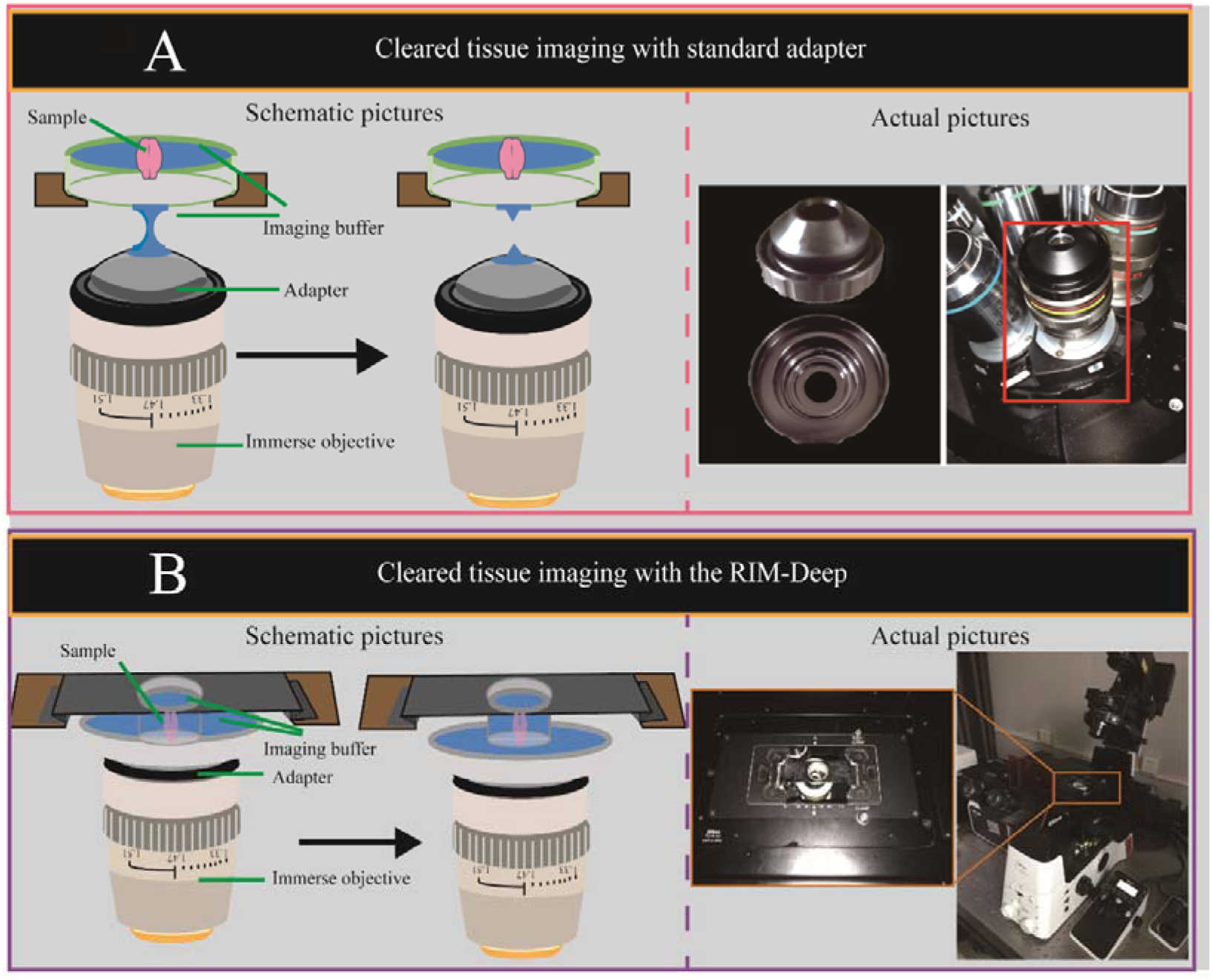
Illustration and actual picture of standard adapter (A) and the RIM-Deep (B) mounted on the Nikon AXR microscope.

**Supplementary figure 3.**
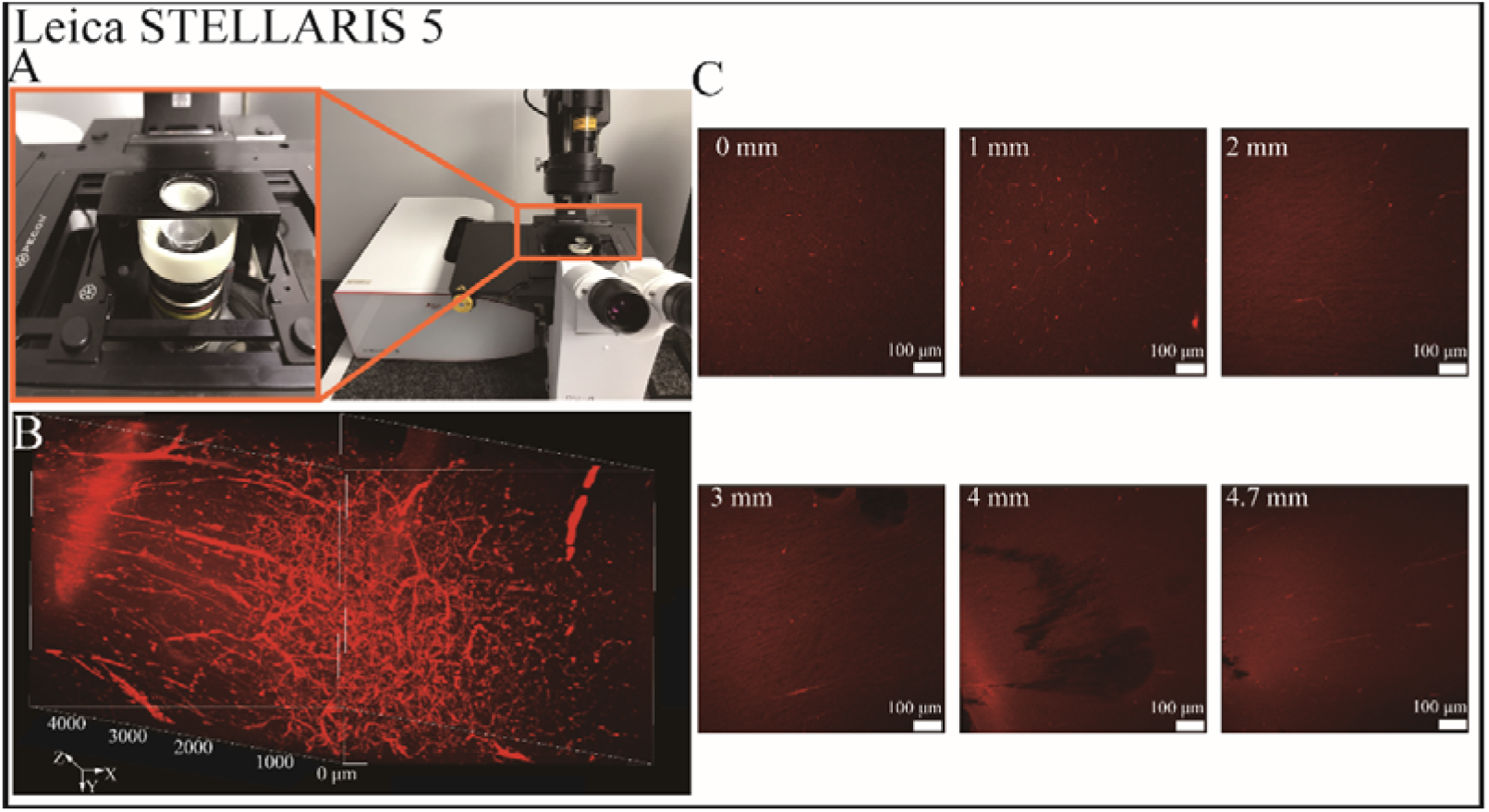
High-depth imaging of *Macaca fascicularis* brain vasculature using Leica STELLARIS 5 with RIM-Deep. (A) Setup of Leica STELLARIS 5 microscope with RIM-Deep assembly. (B) 3D imaging of cleared *Macaca fascicularis* brain vasculature. (C) Optical sections of (B) at varying depths.

